# Life-Stage Transitions Drive Distinct Gut Viral Communities in Dairy Cattle

**DOI:** 10.1101/2025.05.24.655612

**Authors:** Ryan Cook, Adam M. Blanchard, Caleb Marsh, Alise J. Ponsero, Jessica Reynolds, Evelien M. Adriaenssens, Chris Hudson, Jon L. Hobman, Dov J. Stekel, Michael A. Jones, Andrew D. Millard

## Abstract

Ruminant gut microbial communities profoundly influence host health and environmental impacts, yet their viral components remain poorly characterised across developmental transitions. Here, we analysed the dairy cow gut virome across four life stages—calf, heifer, dry adult, and lactating adult. Using hybrid sequencing, we assembled 30,321 viral operational taxonomic units, including 1,338 complete genomes representing mostly novel lineages. Virome composition shifted dramatically with life stage, transitioning from low-diversity, temperate-dominated communities in calves to high-diversity, lytic-dominated communities in adults. Virome transitions paralleled but showed distinct dynamics from bacterial community development, with viral and bacterial diversity negatively correlated during drying-off. Dry cows exhibited elevated viral loads relative to their bacterial hosts. We identified 26 viral sequences targeting the methanogen *Methanobrevibacter*, absent in calves but present in adults. These findings reveal the dynamic nature of ruminant gut viral communities and highlight phages’ potential regulatory role during critical life transitions.

## Main

The farming of cattle constitutes 50% of global livestock units, with an estimated 265 million dairy cows worldwide, underscoring the sector’s pivotal role in food production, agricultural economics, and environmental health^1–3^. Throughout a dairy cow’s life, dramatic physiological and nutritional transitions occur that profoundly reshape gut microbial communities, with likely impacts on health, productivity, and environmental footprint.

Dairy cows experience several distinct developmental stages, each characterised by unique dietary and physiological conditions—from milk-fed calves with developing rumens, to heifers transitioning to high-fiber diets, to lactating and dry adults with cyclical shifts in nutritional requirements ^4^. These transitions create ecological opportunities for microbial community reorganisation and potentially periods of enhanced vulnerability to dysbiosis.

While bacterial community dynamics across these life stages have received increasing attention^5–7^, our understanding of the viral community—the virome—remains limited, particularly for the intestinal tract. Previous work has characterised the virome of agricultural slurry^8^, and examined rumen viromes^9,10^, but comprehensive analyses of intestinal viromes across key life-stage transitions in dairy cattle are lacking.

Viral communities likely play crucial roles in shaping bacterial population dynamics through predation, horizontal gene transfer, and metabolic reprogramming. Bacteriophages can dramatically alter microbial community structure through “Kill-the-Winner” dynamics and “Piggyback-the-Winner” scenarios^11,12^, potentially influencing health outcomes and productivity during stress-associated transition periods in dairy cattle life stages.

Here, we characterised faecal viromes and metagenomes of Holstein-Friesian dairy cows at four key life stages to elucidate how microbial communities develop alongside physiological and nutritional changes. By analysing both components from the same animals, we reveal not only compositional shifts in each domain but also the ecological interactions between viruses and their bacterial hosts throughout dairy cattle development.

## Result

### A Compendium of Complete Viral Genomes from the Dairy Cow Gut

We generated twenty ultra-deep cow gut viromes across four life stages (five each of calves, heifers, dry adults, and lactating adults, Figure 1A) using a combination of long and short read sequencing. Illumina sequencing yielded ∼277 Gb of data (median library sizes: calves 1.7 Gb, heifers 14.9 Gb, dry adults 16.5 Gb, lactating adults 19.4 Gb, overall 14.3 Gb), and a pooled Nanopore library produced ∼104 Gb. Implementation of a short-read exclusion kit for Nanopore sequencing improved median read lengths (6.0-6.9 kb per flow cell) relative to sequencing without exclusion (1.7-3.0 kb per flow cell), without reducing data yields (Supplementary Figure 1). Complementary bacterial metagenomes were generated from the same samples, producing ∼162 Gb of sequence data (median library sizes: calves 8.6 Gb, heifers 7.8 Gb, dry adults 8.1 Gb, lactating adults 7.7 Gb, overall 8.0 Gb).

**Figure 1.**
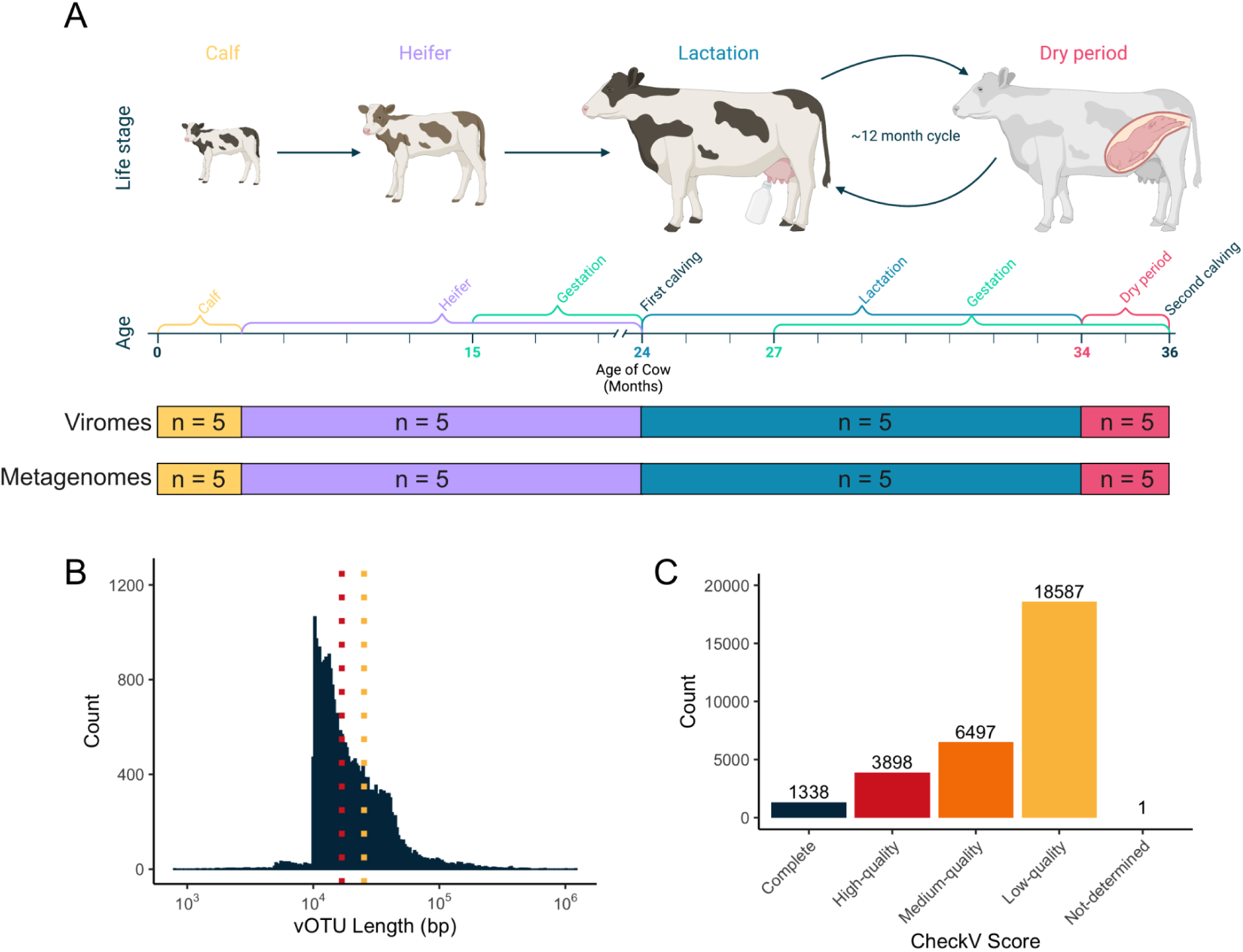
Assembly and quality assessment of viral genomes from dairy cattle faeces. (**A**) Schematic overview of key life stages in a typical UK dairy cow during the first two years of life, corresponding to the sampling framework used in this study. (**B**) Length distribution of viral Operational Taxonomic Units (vOTUs), with median (16.9 kb) and mean (25.3 kb) lengths indicated by vertical intercepts. (**C**) Quality assessment of vOTUs using CheckV.

Viral contigs were assembled, filtered and de-replicated to produce 30,321 unique vOTUs that are approximate to viral species, ranging from 1.5 kb to 1.2 Mb in length with mean and median lengths of 25.3 and 16.9 kb, respectively (Figure 1B). Quality assessment using CheckV classified 1,338 vOTUs as complete, 3,898 as high-quality, 6,497 as medium-quality, 18,587 as low-quality, and one as undetermined (Figure 1C). Manual annotation checks confirmed the absence of non-viral elements within the 1,338 complete genomes.

Taxonomic analysis of the 1,338 complete genomes revealed striking novelty in the bovine gut virome. At the level of Realm, 1,249 complete genomes were classified as *Duplodnaviria* (dsDNA viruses; Figure 2A) and 89 as *Monodnaviria* (ssDNA viruses; Figure 2B). Of these, only 105 complete genomes fell within previously described families—*Intestiviridae* (n=1), *Steigviridae* (n=4), *Salasmaviridae* (n=10), *Microviridae* (n=84), *Autotranscriptviridae* (n=5), and *Peduoviridae* (n=1). The remaining 1,233 complete genomes (92%) did not fall within known families, often forming deeply branching monophyletic clades lacking ICTV-classified members (Figure 2AB), highlighting the exceptional novelty of the bovine gut virome. The distinct clustering patterns and deep branches in the phylogenetic analysis suggest these genomes likely represent numerous novel viral families and potentially new orders awaiting formal taxonomic classification.

**Figure 2.**
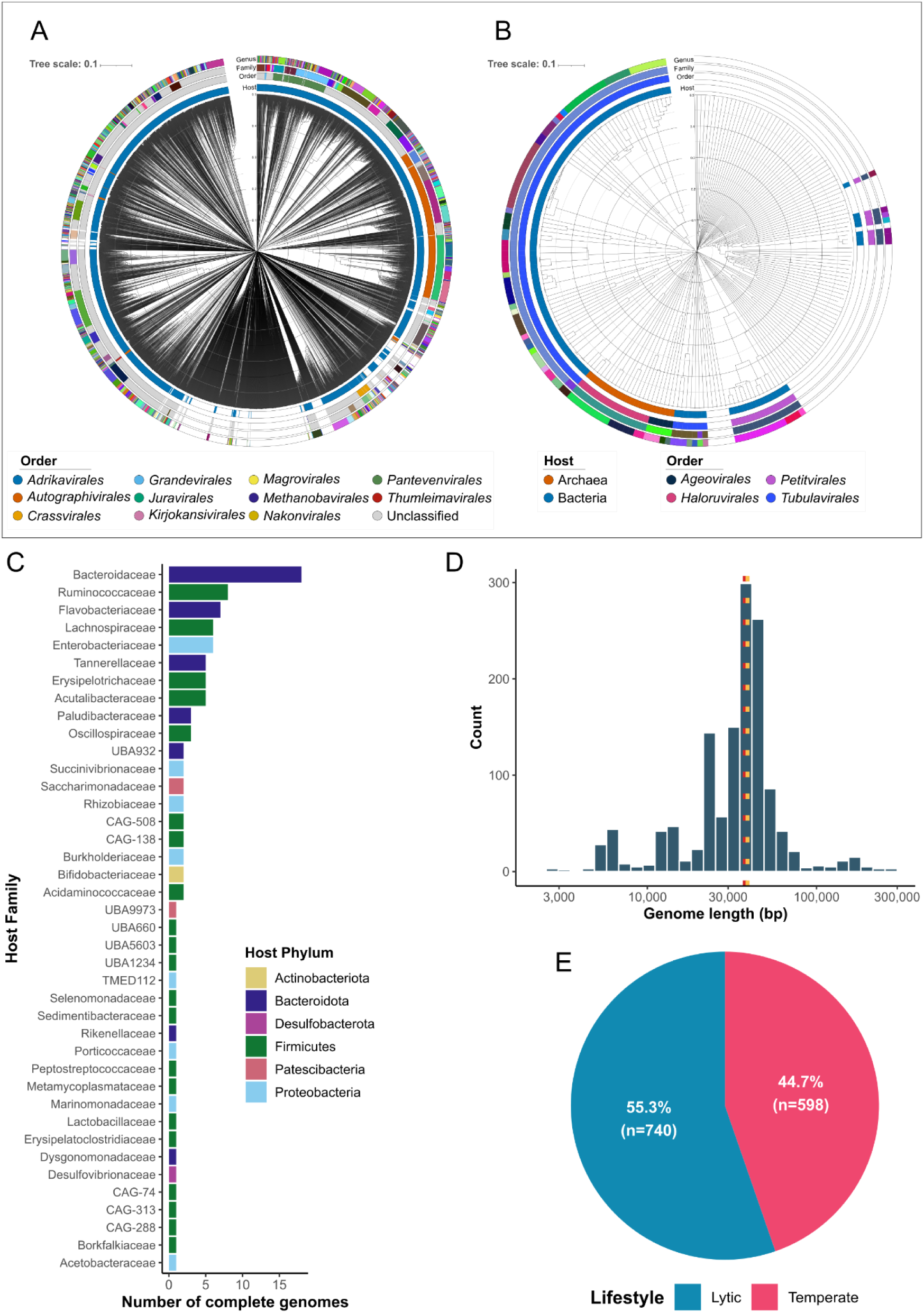
Characteristics of complete viral genomes from the bovine gut virome. (**A**) Proteome-wide phylogenetic comparison of 1,249 complete dsDNA viral genomes (*Duplodnaviria*) from the dairy cow gut with all classified *Duplodnaviria* (VMR40, March 2025). (**B**) Similar to A, for the 89 ssDNA viral genomes (*Monodnaviria*) alongside all classified *Monodnaviria* (VMR40, March 2025). Concentric rings show Host (Bacterial or Archael), Order, Family, and Genus of classified viruses. Legends are omitted for Family and Genus due to the high numbers, with colouring used to differentiate groups. (**C**) Host predictions for complete viral genomes shown at the level of Family and coloured by Phylum. (**D**) Length distribution of complete genomes with mean (39.1 Kb) and median (37.8 Kb) indicated by yellow and red dashed lines respectively. (**E**) Proportion of complete viral genomes predicted to be temperate and lytic.

Lifestyle prediction indicated that 598 complete genomes (44.7%) were likely temperate phages capable of integration into host genomes, while the remaining 740 (55.3%) were predicted to be obligately lytic (Figure 2E). Host prediction was possible for only 105 of the complete genomes, with the vast majority (1,233) lacking a confident host assignment (Figure 2C). Among those with predicted hosts, bacteria classified within the *Firmicutes* were the most commonly targeted phylum (n=46), followed by *Bacteroidota* (n=37), *Proteobacteria* (n=16), and *Patescibacteria* (n=3). At the family level, viruses targeting bacteria within the Bacteroidaceae (n=18), Ruminococcaceae (n=8), and Enterobacteriaceae (n=6) were most frequent, consistent with the prevalence of these bacterial families in the ruminant gut^5^. Notably, many of the predicted hosts represent important members of the cow gut microbiota, including *Faecalibacterium* (n=4), *Bacteroides* (n=11), *Prevotella* (n=7), and *Escherichia* (n=4).

### Life-Stage Transitions Drive Distinct Viral Community Profiles

Of the 20 faecal viromes sequenced, two calf samples did not meet our vOTU detection threshold (≥1× coverage over 75% of vOTU length) and were excluded, leaving 18 viromes for downstream analyses. Parallel bacterial metagenomic data were available for these samples, allowing us to examine both bacterial and viral community dynamics across the four major life stages.

Alpha diversity analyses revealed stark differences between calves and adults for both viral and bacterial communities. Calf viromes exhibited markedly lower Shannon diversity (mean 2.49 ± 0.45) compared to heifers (7.33 ± 0.52), dry adults (7.68 ± 0.16), and lactating adults (6.58 ± 1.97) (Kruskal-Wallis test, χ² = 12.68, p = 0.005; Figure 3A). This pattern was even more pronounced when examining community evenness through Inverse Simpson index, with calves showing extremely low evenness values (mean 5.73 ± 1.40) compared to the substantially more balanced viral communities in heifers (270.63 ± 105.51), dry adults (317.80 ± 85.03), and lactating adults (130.77 ± 98.85) (Kruskal-Wallis test, χ² = 12.15, p = 0.007; Figure S2). Post-hoc pairwise comparisons using Dunn’s test with Bonferroni correction confirmed significant differences in evenness between calves and each adult group (p < 0.01 for all comparisons), while differences among adult groups were not statistically significant (p > 0.05), despite the notably higher evenness in dry adults. The dramatic >50-fold increase in evenness from calves to adults indicates not just greater viral richness but a fundamental restructuring towards communities where abundance is more equitably distributed across viral taxa. This pattern was mirrored in nucleotide-level variation, with adults displaying substantially greater micro-diversity than calves (Figure 3B).

**Figure 3.**
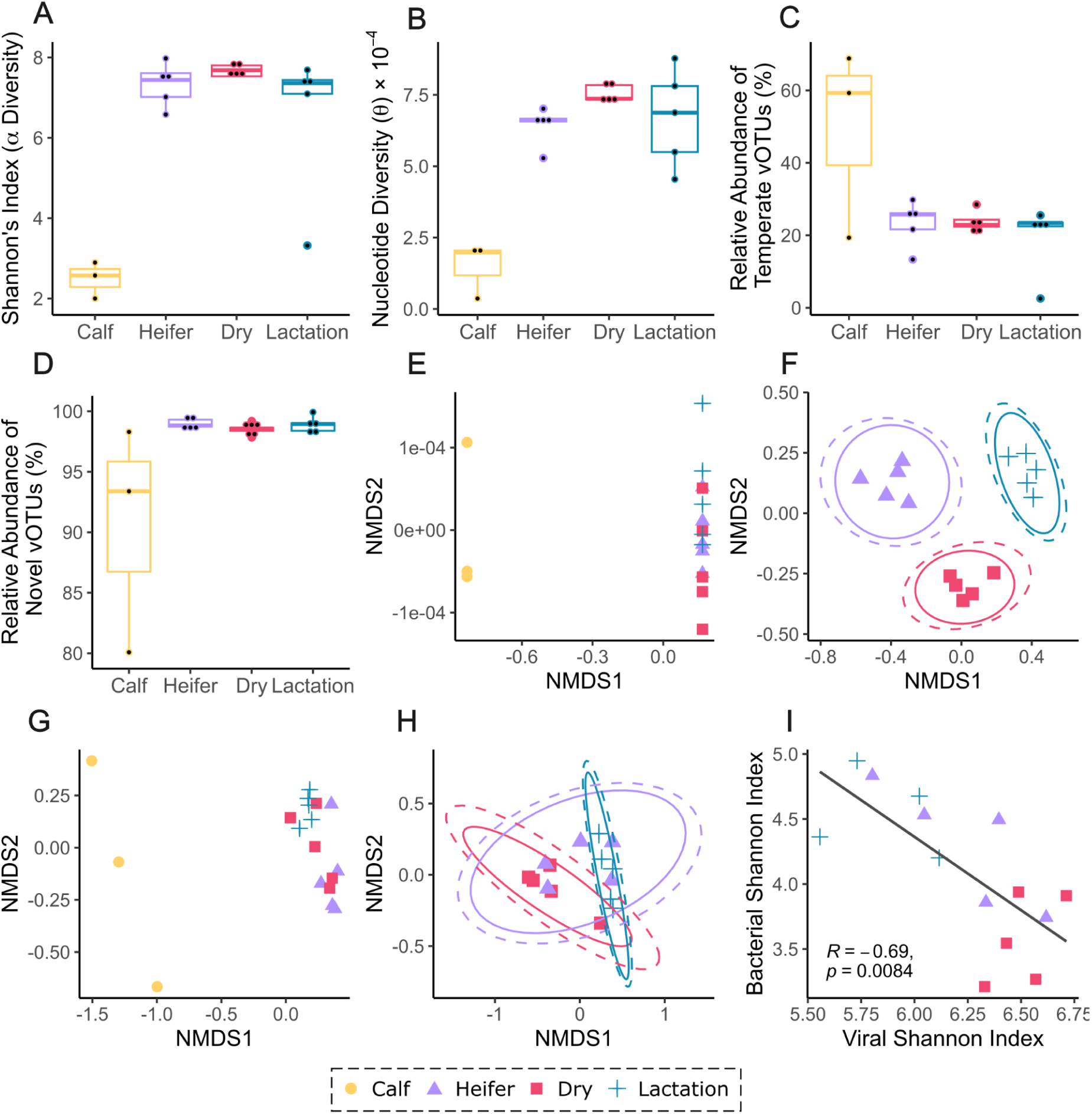
Compositional shifts and diversity patterns of viral and bacterial communities across dairy cattle life stages. (**A**) Shannon’s alpha diversity index for viral communities across four life stages (Kruskal-Wallis test, χ² = 12.68, p = 0.005). (**B**) Nucleotide microdiversity (theta) showing within-sample genetic variation. (**C**) Relative abundance of temperate vOTUs across life stages (Kruskal-Wallis test, χ² = 1.69, df = 3, p = 0.639). (**D**) Relative abundance of novel vOTUs not assigned to known genera (Kruskal-Wallis test, χ² = 7.31, df = 3, p = 0.063). Beta diversity analysis based on Bray-Curtis dissimilarity of the dairy cow virome shown as PCoA plots (**E**) including all samples (PERMANOVA, R² = 0.10, P = 0.001) and (**F**) excluding calves to better visualise differences among adult stages (PERMANOVA, R² = 0.17, P = 0.001). Similar Bray-Curtis dissimilarity analysis of bacterial communities (**G**) including all samples (PERMANOVA, R² = 0.48, P = 0.001) and (**H**) excluding calves (PERMANOVA, R² = 0.15, P = 0.023). (**I**) Negative correlation between bacterial and viral diversity measured by Shannon’s index (Spearman’s rho = –0.69, P=0.0084).

The bacterial communities showed similar life-stage stratification, though with an intriguing inverse relationship to viral diversity—while viral diversity increased during the transition from lactating to dry adult status, bacterial diversity decreased. This negative correlation between bacterial and viral diversity (Spearman’s rho = –0.69, P=0.0084) was particularly pronounced during the drying-off period (Figure 3I), suggesting potential ecological interactions between the two communities.

Beta-diversity analyses reinforced the distinctiveness of life-stage communities in both viral and bacterial domains. Using Bray-Curtis dissimilarity, both viral and bacterial communities showed significant clustering by life stage in nested PERMANOVA tests (999 permutations). For viral communities, the primary division between calves and adults explained 10.3% of the total variation (R² = 0.10, P = 0.001), while differences among the three adult stages (heifers, dry, and lactating) explained an additional 17.4% of variation (R² = 0.17, P = 0.001; Figure 3EF). Bacterial communities demonstrated an even stronger age-related structuring, with the calf-adult division explaining 47.8% of total variation (R² = 0.48, P = 0.001)—more than four times the proportion explained in the viral dataset. Differences among adult stages contributed an additional 15.1% of bacterial community variation (R² = 0.15, P = 0.023; Figure 3GH).

Testing for homogeneity of multivariate dispersions confirmed that these differences were driven by distinct community centroids rather than within-group dispersion effects for both viral (permutest, P = 0.352) and bacterial communities (permutest, P = 0.554). The strong association between viral and bacterial community structures was confirmed by both Mantel test (r = 0.671, P = 0.001) and Procrustes analysis (correlation = 0.740, P = 0.003), indicating coordinated ecological shifts in both domains across life stages. The compositional changes in viral communities closely parallel those in bacterial communities, suggesting interconnected ecological dynamics between these two components of the bovine gut ecosystem.

### Ecological Shift from Temperate to Lytic Viral Communities During Development

The composition of viral communities differed dramatically between calves and adults. Calf viromes contained a higher proportion of temperate phages (49.15 ± 26.41%) compared to heifers (23.32 ± 6.40%), dry adults (23.67 ± 3.13%), and lactating adults (19.48 ± 9.33%), though this difference did not reach statistical significance (Kruskal-Wallis test, χ² = 1.69, df = 3, p = 0.639; Figure 3C). This suggests a potential shift toward predominantly lytic viral communities in adults (>76% of viral abundance), which may represent an ecological transition during ruminant development, though larger sample sizes would be needed to confirm this pattern statistically.

Taxonomic assignment alongside known viruses further supported this developmental transition. The relative abundance of vOTUs not assigned to known genera was lowest in calves (90.59 ± 9.33%) compared to adult animals, with heifers (98.99 ± 0.53%), dry adults (98.53 ± 0.50%), and lactating adults (98.90 ± 0.71%) showing similar levels of novel viral abundance (Kruskal-Wallis test, χ² = 7.31, df = 3, p = 0.063; Figure 3D). Public database mapping patterns supported these findings, with calves averaging a substantially higher fraction of reads mapping to known phages (Figure 4AF).

**Figure 4.**
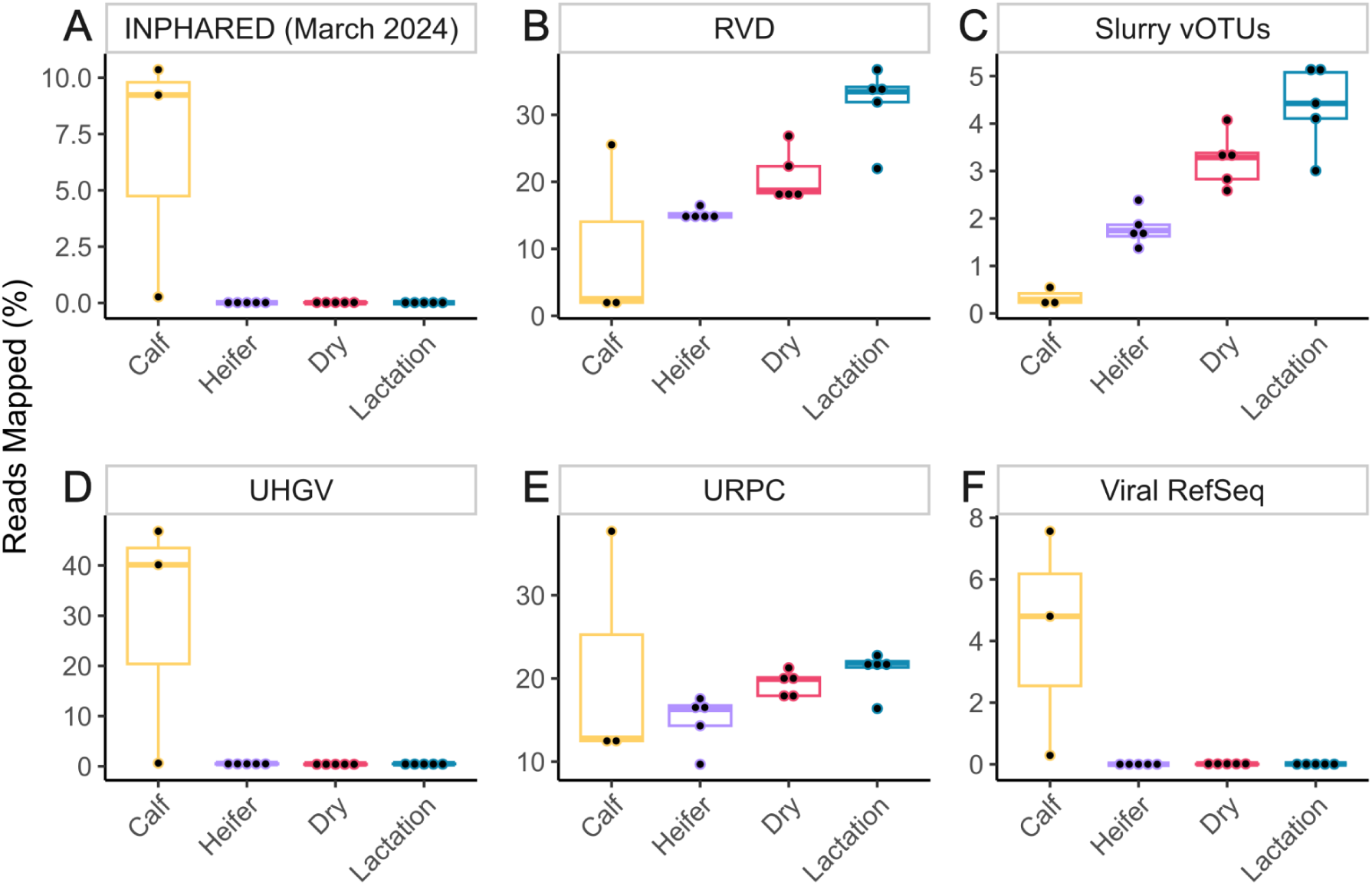
Mapping patterns of viral reads to reference databases across dairy cattle life stages. Percentage of reads from each life stage mapping to different viral databases: (**A**) INPHARED (March 2024; Kruskal-Wallis test, χ² = 7.89, p = 0.048), (**B**) Rumen Virome Database (RVD; Yan et al. 2023; χ² = 10.73, p = 0.013), (**C**) farm-specific dairy slurry virome (Cook et al. 2021; χ² = 10.17, p = 0.017), (**D**) Unified Human Gut Virome Catalog (UHGV; χ² = 8.63, p = 0.035), (**E**) Unified Rumen Phage Catalogue (URPC; Wu et al. 2024; χ² = 7.62, p = 0.049), and (**F**) NCBI Viral RefSeq (March 2024; χ² = 8.05, p = 0.045).

Comparison with existing viral databases further confirmed the life-stage stratification of viral communities. Mapping rates varied significantly across life stages for human gut viral databases (Kruskal-Wallis test, χ² = 8.63, p = 0.035; Figure 4), with calf viromes exhibiting remarkably higher mapping frequency (UHGV: 0.63-46.81%) compared to adult cattle (0.3-0.63%). Similarly, calves showed significantly higher mapping to general viral reference collections (INPHARED: 0.27-10.36%, χ² = 7.89, p = 0.048; Viral RefSeq: 0.29-7.56%, χ² = 8.05, p = 0.045; Figure 4) compared to adult cattle (<0.03% for both databases), suggesting that calf viromes contain more similarity to previously characterised viruses.

Conversely, the slurry tank virome from the same farm showed significantly higher mapping from adult samples (Kruskal-Wallis test, χ² = 10.17, p = 0.017; Figure 4), with highest similarity to lactating (3.01-5.19%) and dry cows (2.59-4.07%), followed by heifers (1.37-2.39%), with minimal mapping from calves (0.16-0.55%). This gradient reflects the population structure of the farm, where lactating cows comprise the largest proportion of animals and therefore contribute most significantly to the slurry tank microbiome. All adult stages also showed moderate mapping to ruminant-specific viral databases, with both RVD (14.38-36.74%, χ² = 10.73, p = 0.013; Figure 4) and URPC (9.68-22.76%, χ² = 7.62, p = 0.049; Figure 4) showing significant differences across life stages. Lactating cows exhibited particularly high mapping rates to the RVD database (21.98-36.74%). These mapping patterns align with the observed ecological shifts in viral communities during development and further support the transition from a calf virome with human gut-like characteristics to a specialised ruminant virome in adults.

The core virome analysis further highlighted the developmental transition in viral communities. After rarefaction, only 225 vOTUs were present in at least one calf sample, of which 47 were detected in any adult group. In contrast, each adult group displayed a substantial core virome—3,385 (heifers), 3,049 (dry adults), and 3,254 (milking adults) vOTUs detected in ≥4/5 individuals. Despite this richness, only 711 vOTUs were shared across all three adult groups, indicating that while adult cows share a considerable viral component, each life stage maintains a partly distinct community.

### Virus-Host Dynamics Reveal Complex Ecological Interactions Across Life Stages

To explore the relationship between viruses and their bacterial hosts across development, we predicted host genera for each vOTU and calculated a virus-host ratio (VHR) by dividing the total relative abundance of viruses by the relative abundance of their predicted microbial hosts^13^. A consistent negative correlation between VHR and host abundance was observed across all adult life stages (Spearman’s rho from -0.61 to -0.74, p < 2.2e-16; Figure 5A). This inverse relationship suggests that more abundant bacterial taxa tended to have fewer associated phages per host genome, consistent with the ecological model of “Kill-the-Winner“^12^ wherein viral predation selectively targets the most abundant bacterial populations, thereby maintaining diversity in the microbial community.

**Figure 5.**
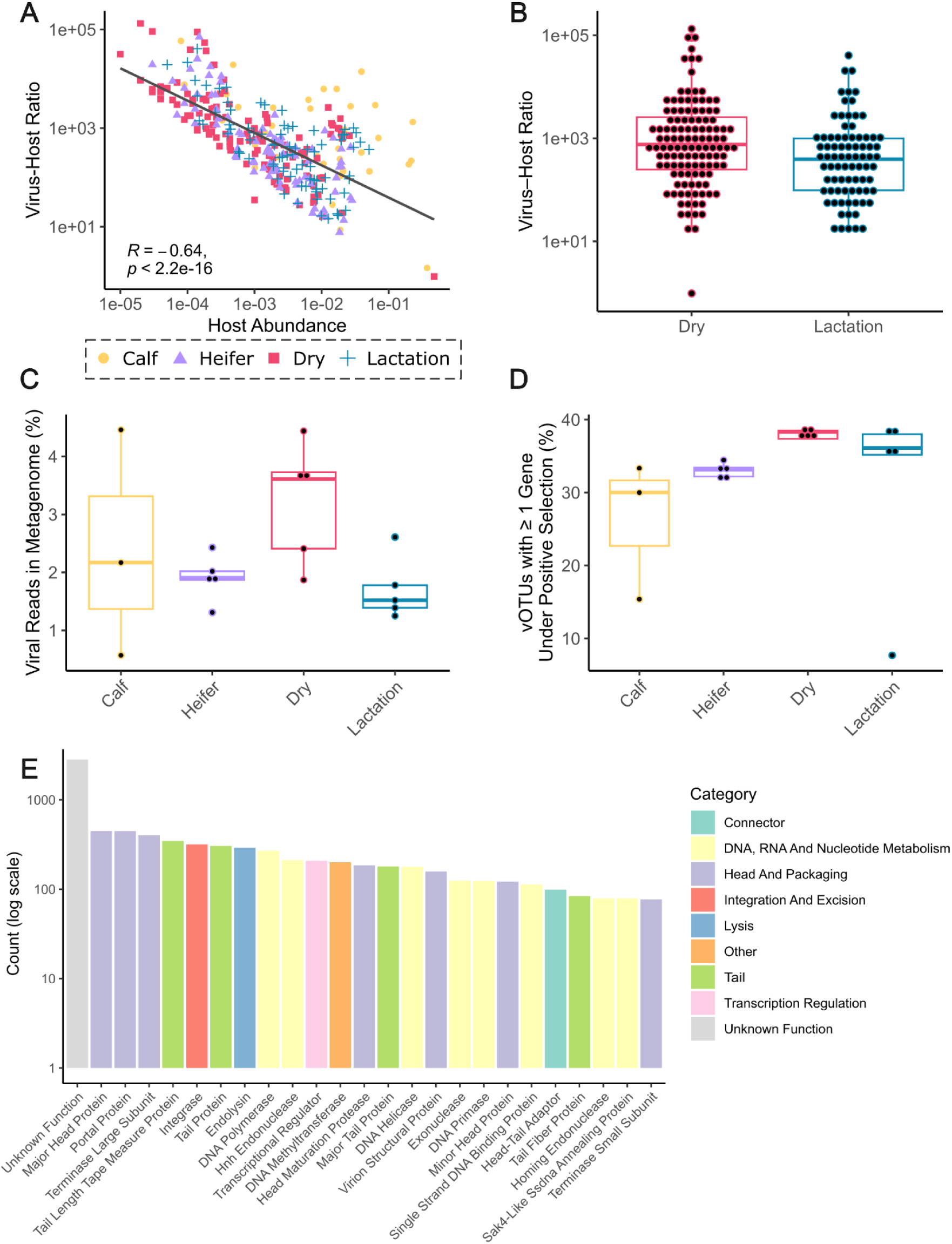
Virus-host ecological dynamics reveal distinct patterns across dairy cattle life stages. (**A**) Inverse relationship between virus-host ratio and bacterial host abundance across all adult life stages (Spearman’s rho from -0.61 to -0.74, p < 2.2e-16), consistent with “Kill-the-Winner” dynamics. (**B**) Comparison of virus-host ratios between dry and lactating adults showing significantly elevated viral loads relative to bacterial hosts during the drying-off period (Mann-Whitney U test, P=0.016). (**C**) Viral load measured as the proportion of microbial metagenomes mapping to vOTUs, demonstrating heightened viral presence in dry adults. (**D**) Percentage of vOTUs with at least one gene under positive selection across life stages, showing significant differences between groups (Kruskal-Wallis test, χ² = 9.73, df = 3, p = 0.021). (**E**) Top 25 most common PHROG annotations of genes under positive selection in at least one sample.

The VHR analysis highlighted the dry cow period as particularly dynamic for virus-host interactions. Dry cows exhibited a significantly higher median VHR compared with lactating adults (Wilcoxon rank-sum test, P=0.006; Figure 5B), indicating a shift toward increased viral loads relative to their bacterial hosts during this critical transition period. To further examine this pattern, we determined viral load by mapping bulk metagenomic reads to our vOTUs, calculating the percentage of total metagenomic DNA attributable to viral sequences. This revealed consistent group differences, with dry cows exhibiting the highest median viral load (3.61% of metagenome), followed by calves (2.17%), heifers (1.91%), and lactating cows showing the lowest (1.52%; Figure 5C). The significant difference between dry and lactating cows (Mann-Whitney U test, P=0.016; Figure 5C) corroborates the VHR analysis and further illustrates heightened viral activity during the drying-off period.

Evidence of ongoing evolutionary dynamics between phages and their bacterial hosts was found across all life stages. A substantial fraction of detected vOTUs in all groups contained at least one gene under positive selection (Figure 5D), with significant differences observed among life stages (Kruskal-Wallis test, χ² = 9.73, df = 3, p = 0.021). Dry adults exhibited the highest proportion of vOTUs under selection (mean 38.0% ± 0.65%), significantly higher than calves (26.2% ± 9.55%; Dunn’s test with Bonferroni correction, p = 0.0197) and marginally higher than heifers (33.0% ± 1.01%, p = 0.0455). Lactating adults showed intermediate levels (31.2% ± 13.2%) that did not significantly differ from other groups. The consistently elevated proportion of vOTUs under selection in dry adults suggests that selective pressures on viral genes are particularly intense during the drying-off period, likely reflecting rapid adaptation to changing host populations or environmental conditions.

Among genes under positive selection, functions related to DNA, RNA and nucleotide metabolism were particularly prominent (2,545 instances; Figure 5E), including DNA polymerase (270), HNH endonuclease (212), and DNA helicase (178). Structural components were also frequently under selection, with head and packaging genes (2,127 instances; Figure 5E) such as major head protein (449) and portal protein (447), as well as tail-related genes (1,270 instances; Figure 5E) including tail length tape measure protein (346) and tail protein (305). The prevalence of selection on phage structural genes, particularly those involved in host recognition and infection, suggests ongoing evolutionary arms races between phages and their bacterial hosts across all life stages, with particularly intense selective pressure during the physiologically critical drying-off period.

### Identification of Archaeal-Targeting vOTUs and Methanogen-Associated AMGs

Beyond bacterial viruses, our analysis also identified viruses targeting archaeal hosts, in particular methanogens important to ruminant digestion^14^. In total, 60 vOTUs were predicted to infect archaeal hosts, including 10 with CRISPR matches to archaeal MAGs from a rumen MAG compendium (table S4)^15^. Of these, 26 vOTUS were inferred to target *Methanobrevibacter*, the primary genus of methanogens in the bovine gut (Figure 6A). These putative *Methanobrevibacter*-infecting vOTUs ranged from 10 to 470 Kb in length, with seven estimated to be >90% complete by CheckV. Seven were predicted to be temperate, and the remaining 19 could not be assigned a lifestyle, potentially being obligately lytic. Taxonomic classification assigned all to *Caudoviricetes*, with six belonging to the Order *Methanobavirales* (Figure 6C). The remaining 20 likely represent seven novel orders of tailed archaeal viruses that contain no cultivated members (Figure 6C).

**Figure 6.**
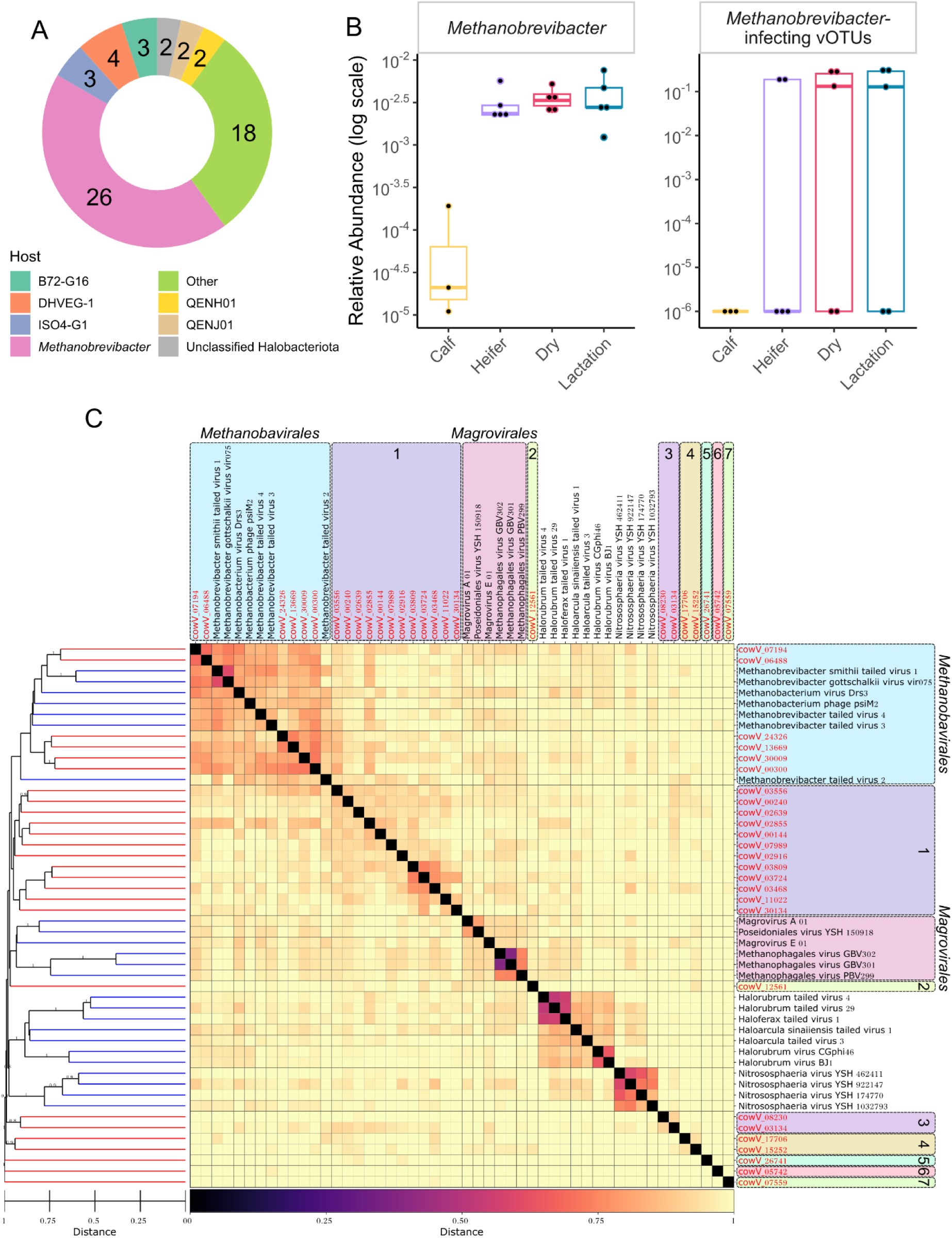
Methanogen-targeting viruses emerge during ruminant development with potential implications for methane regulation. (**A**) Distribution of vOTUs predicted to infect different archaeal genera. (**B**) Relative abundance of *Methanobrevibacter* and its predicted viruses across life stages. (**C**) Proteome-based clustering analysis of *Methanobrevibacter*-targeting vOTUs in relation to classified tailed archaeal viruses within *Caudoviricetes*, revealing seven potential novel viral orders lacking cultivated representatives. Distances between pairs of genomes are calculated as a composite generalised Jaccard distance of protein profile HMM similarity and genome organisation.

The archaeal viruses showed distinct life-stage patterns similar to bacterial viruses. None of the *Methanobrevibacter*-targeting vOTUs were detected in calf samples but appeared in over half of the adult samples, albeit at low absolute abundance (0.01–0.06% relative abundance when present; Figure 6B). This pattern paralleled the dramatic expansion of *Methanobrevibacter* itself, with its relative abundance being 75,600% higher in adults (0.00123-0.00757%; Figure 6B) compared to calves (0.00001-0.00019%; Figure 6B). This co-development of methanogen populations and their associated viruses suggests a coordinated ecological succession as the rumen matures. The complete absence of methanogen viruses in calves, followed by their consistent presence in adult ruminants, indicates that viral regulation of methane-producing archaea emerges as a functional component of the mature ruminant gut ecosystem.

The potential impact of these viruses on methanogen metabolism is further supported by the presence of several putative auxiliary metabolic genes (AMGs) associated with methane pathways (KEGG map00680). The most frequently identified—predicted in 49 vOTUs—was glycine hydroxymethyltransferase (glyA/SHMT; K00600) that is involved in one-carbon metabolism, and can play a role in the production of methionine. Methionine can act as a source of methyl groups that may form methane. More directly relevant to methanogen metabolism, we detected two vOTUs containing coenzyme F420 hydrogenase subunit beta (frhB; K00441), one vOTU with heterodisulfide reductase subunit A2 (hdrA2; K03388), and one vOTU encoding gamma-F420-2 ligase (cofF; K14940). Additional metabolic genes in the map00680 pathway included phosphate acetyltransferase (pta; K00625, two vOTUs), 6-phospho-3-hexuloisomerase (hxlB; K08094, two vOTUs), glycerate 2-kinase (gck; K11529, one vOTU), and phosphofructokinase (pfk/pfp; K21071, one vOTU). Taken together, these findings suggest viruses may interact with methanogen populations through both direct predation and metabolic modulation in the developing bovine gut ecosystem.

## Discussion

The viral community of dairy cows has been largely underexplored. Previous work has focused on specific stages of a lifecycle (e.g., lactation) where very few complete bacteriophages were identified ^16^, or has been limited to characterising viruses within the rumen ^9,17,18^. Here we utilised a combination of short and long read sequencing to gain a comprehensive understanding of gut viral diversity in dairy cattle across key developmental stages, identifying >30,000 vOTUs, including 1,338 complete and 3,898 high-quality viral genomes.

We observed a stark contrast between calf and adult viromes. Calves harboured significantly lower viral diversity, with communities dominated by recognisable, often temperate phages—largely sharing similarity with human gut viral sequences. The dominance of temperate phages in calves may result from induction of prophages from early colonising bacteria, as has been proposed in human infant gut development^19,20^.

As the gastrointestinal tract matures and the diet diversifies post-weaning, the virome undergoes a dramatic expansion in both diversity and complexity. The adult virome was characterised by high novelty, with the vast majority of complete viral genomes representing previously undescribed viral lineages. This pattern mirrors developmental progressions observed in human infant gut viromes^21–24^. However, unlike human gut viromes that typically stabilise in adulthood, the dairy cow virome continues to undergo significant compositional shifts across adult life stages, reflecting the distinct physiological transitions these animals experience throughout their productive life.

The shift from temperate to lytic viral communities with age likely reflects changing ecological strategies as the gut ecosystem matures. The predominance of temperate phages in calves may provide evolutionary advantages in a developing, less stable environment, while the transition to lytic dominance in adults could indicate a shift toward competition-based dynamics in a more stable ecosystem, consistent with the “Kill-the-Winner” model of viral-bacterial coevolution.

Our integration of bacterial and viral metagenomic data revealed complex ecological relationships between these communities. The strong correlation between bacterial and viral beta diversity confirmed that these communities largely shift in concert across life stages. However, the negative correlation between bacterial and viral alpha diversity—particularly pronounced during the drying-off period—suggests a more nuanced relationship than simple co-development.

The drying-off period emerged as a particularly dynamic phase for virus-host interactions. The transition involves substantial dietary changes, with reduced energy intake and a shift toward more fibrous rations. These changes coincided with increased viral diversity, decreased bacterial diversity, and a significantly higher virus-host ratio. Furthermore, dry cows exhibited the highest proportion of viral genes under positive selection, suggesting intensified evolutionary dynamics between phages and their bacterial hosts during this transition. Our identification of 26 vOTUs targeting *Methanobrevibacter*, absent in calves but widely present in adult samples, provides temporal context to emerging research on archaeal viruses in the rumen ecosystem^17,25^. The life-stage specificity suggests archaeal viruses emerge alongside the establishment of functional methanogen populations during rumen development, which may inform future strategies for methane mitigation through virome manipulation.

From an applied perspective, our findings have potential implications for dairy cattle management. The identification of critical transition periods characterised by viral community restructuring—particularly weaning and drying-off—suggests these phases may represent intervention opportunities for modulating the microbiome. Future research should explore the functional implications of these virome transitions, particularly the role of phage-mediated gene transfer in conferring adaptive traits to bacterial hosts across developmental stages.

In conclusion, our work establishes a comprehensive foundation for understanding viral community development in dairy cattle, revealing that life-stage transitions drive profound shifts in virome composition, diversity, and virus-host dynamics. These findings highlight the dynamic nature of phage-bacteria relationships in the ruminant gut and underscore the potential importance of viral communities in shaping microbial ecosystem function across the life cycle of dairy cattle.

## Online Methods

### Sample Collection and VLP Enrichment

Cow faeces was collected from a UK dairy farm using a drop catch method (i.e., catching the sample in a sterile tube before it can contact the ground). Samples (n = 20) were collected from five; pre-weaning calves (< 30 days old), heifers, dry adults, and lactating adults. Samples were kept on ice and processed the same day. Sample collection for this project was reviewed and approved by the SVMS ethics committee on the 14th of November 2017 with approval number 2132 171010.

Viral-like particle (VLP) enrichment was performed as described previously ^8^. Briefly, faeces was homogenised with PBS buffer and centrifuged, prior to filtration to remove bacteria. Viral particles were concentrated using an Amicon column (Sigma-Aldrich) and DNA was extracted using a standard phenol-chloroform extraction.

### Bacterial DNA Extraction and Sequencing

Bacterial DNA was extracted using QIAamp PowerFecal Pro DNA Kit (Qiagen) following manufacturer’s instructions. Individual samples were sequenced using Illumina (2 X 150 bp) short-read sequencing with NextSeq 2000 at the Institute of Molecular Medicine, University of Leeds.

### Virome Short-Read Sequencing

All 20 samples were independently sequenced using Illumina (2 X 150 bp) short-read sequencing by NUomics at Northumbria University. DNA was quantified using Qubit high sensitivity and normalised to 2 ng per library starting concentration. The libraries were prepared using DNAprep (M) (Illumina, San Diego, CA, USA) with unique dual indexes as per manufacturer’s instructions. The library was checked using BioAnalyzer (Agilent, Santa Clara, CA, USA) and Qubit high sensitivity and normalised to 30 nM. The libraries were pooled and ran on a MiSeq V2 300 cycle nano kit prior to sequencing on the Novaseq 6000 300 cycle SP kit.

### Virome Long-Read Sequencing

DNA from the 20 samples was pooled and amplified using the REPLI-g Mini kit following the manufacturer’s instructions (Qiagen, Valencia, CA, USA) to gain sufficient material for ONT sequencing. Amplified DNA was de-branched using S1 Nuclease (Promega) at 10 units per μg of DNA (quantified using a Qubit) to minimise chimeras introduced during amplification^26^, followed by passage through a Zymo Clean & Concentrator-25 column. To enrich for high molecular weight DNA, a 10 kb short read exclusion kit (Circulomics, Baltimore, MD, USA) was used following the manufacturer’s instructions, with the following modifications. The DNA pellet was resuspended in 50 μl of nuclease-free water rather than the provided buffer. Libraries were prepared using the SQK LSK-110 ligation sequencing kit (ONT, Oxford, UK) prior to sequencing on ten MinION flow cells (six r9.4.1 and four r10.3), with four out of ten being loaded with DNA that had been processed using the short read exclusion kit prior to library preparation.

### Virome Quality Control and Assembly

Adapters were trimmed from short reads using bbduk.sh v38.84 with ktrim=r minlen=40 minlenfraction=0.6 mink=11 tbo tpe k=23 hdist=1 hdist2=1 ftm=5 ref= /bbmap/resources/adapters.fa^27^ followed by quality trimming with maq=8 maxns=1 minlen=40 minlenfraction=0.6 k=27 hdist=1 trimq=12 qtrim=rl^27^ . Tadpole.sh v38.84 was used to correct sequencing errors with mode=correct ecc=t prefilter=2^27^ . Trimmed reads were mapped to the *Bos taurus* reference genome (NKLS00000000) using bbmap.sh v38.84 with local=t minid=0.95 maxindel=6 tipsearch=4 bandwidth=18 bandwidthratio=0.18 usemodulo=t printunmappedcount=t idtag=t minhits=1, and unmapped reads were split back into paired end files using reformat.sh v38.84)^27^. VLP enrichment of samples was estimated using ViromeQC v1.0^28^. Library sizes were calculated using the count command within SeqFu v1.22.3 ^29^. Assembly was performed using MEGAHIT v1.1.1-2-g02102e1 with ‘--k-min 21 --k-max 149 --k-step 24’ and contigs ≥ 1.5 kb were retained^30^.

Long reads were pooled, and low-quality reads removed using Filtlong v0.2.1 with --min_length 1000 --keep_percent 95 (https://github.com/rrwick/Filtlong). Assembly was performed using Flye v2.8.1-b1676 with --meta --min-overlap 1000^31^. Long read polishing was performed with Medaka v1.6.0 with -b 50 (https://github.com/nanoporetech/medaka) in two rounds, first with reads obtained from r9.4.1 flow cells followed by reads obtained from r10.3 flow cells. Illumina reads were pooled, and forward and reverse reads were mapped separately to the medaka-polished assembly using bwa v0.7.12-r1039 to generate SAM files (e.g., bwa mem -a medaka_polished.fa pooled_R1.fq.gz > alignments_1.sam and bwa mem -a medaka_polished.fa pooled_R2.fq.gz > alignments_2.sam)^32^. Alignments were filtered using polypolish_insert_filter.py and the medaka-polished contigs were polished using Polypolish v0.5.0^33^. The polished ONT contigs were processed using CheckV v0.9.0 and any with a kmer frequency of ≥ 1.5 were excluded to remove potential chimeras^34^.

### Filtering vMAGs and vOTUs

The 20 samples of Illumina reads were mapped separately to the Illumina virome assembly using minimap2 v2.17-r941 with -ax sr and sorted BAM files were produced using samtools v1.9^32^. The Illumina assembly and BAM files were used as input for vRhyme v1.1.0 to produce vMAGs^35^. vMAGs from the “best bins” were concatenated into single contigs padded with N’s using concatenate.sh v38.84 and processed using CheckV v0.9.0^27,34^. Bins that obtained a CheckV quality estimate of “low-quality” or “not-determined”, a protein redundancy > 1, a contamination estimate > 10%, the warning flag “no viral genes”, or ≥ 25% “host” genes (and not identified as a prophage) were excluded from filtering. Bins ≥ 10 kb (and one < 10 kb bin that was estimated to be complete due to a high confidence DTR) that satisfied at least one of the following conditions were included: (1) predicted viral by VIBRANT v1.2.0^36^), (2) obtained an adjusted P-value from DeepVirFinder of ≤ 0.05^37^, or (3) had a significant (-E 0.001) to either the viral RefSeq or INPHARED databases using MASH v2.0 (July 2022)^38,39^. For any tool used to process the concatenated vMAGs that uses Prodigal to predict open reading frames (ORFs), the -m flag was manually added to their code so ORFs were not predicted over ambiguous bases (Ns).

Illumina and polished ONT contigs ≥ 10kb and those predicted to be circular (determined using apc.pl (https://github.com/jfass/apc)) were de-replicated using the MIUVIG recommended parameters (95% ANI over 85% length of the shorter sequence) with blast and CheckV scripts as described in the CheckV documentation (https://bitbucket.org/berkeleylab/checkv/src/master/)^34,40,41^. Contig clusters that belonged to an included vMAG were excluded from further analysis. The remaining clustered contigs were filtered using the same three criteria as the vMAGs. The filtered contigs were processed using CheckV v0.9.0 and those with the “no viral genes” warning, < 3 total genes, or ≥ 25% “host” genes (and not identified as a prophage) were excluded^34^. For those identified as prophages, the CheckV trimmed versions were used in downstream analyses ^34^ The contigs and vMAGs that passed filtering formed the 30,321 vOTUs included in this analysis.

The annotations of vOTUs predicted complete by CheckV (n = 1,338) were manually inspected to determine if the vOTU was demonstrably viral (e.g., presence of viral signature genes (such as terminase, portal, tail, capsid etc), a high number of hypothetical proteins, and few genes typically associated with bacteria).

### Functional Annotation and AMG Identification

vOTUs were annotated using Prokka v1.14.6 with a publicly available set of HMMs derived from PHROGs (http://s3.climb.ac.uk/ADM_share/all_phrogs.hmm.gz)^42,43^ . Putative AMGs were identified in the VIBRANT output^36^.

### Taxonomic Assignments

All vOTUs were processed alongside the INPHARED database (August 2022)^39^ using vConTACT2 with --rel-mode ‘Diamond’ --db ‘None’ --pcs-mode MCL --vcs-mode ClusterONE --min-size 1^44^. If a vOTU was in the same VC as a reference sequence, it was considered to belong to known genera, with the remaining considered novel. Predicted complete phage genomes were processed using ViPTreeGen v1.1.3 alongside all *Monodnavira* and *Duplodnavira* that infect bacteria and archaea in the VMR40 (March 2025) and the resulting trees were visualised using IToL^45,46^. The vOTUs predicted to infect *Methanobrevibacter* were processed alongside all archaeal *Caudoviricetes* using GRAViTy-V2 v2.2 using default settings for the new_classification_full pipeline^47^.

### Lifestyle and Host Prediction

vOTUs that may be able to access a lysogenic lifestyle (described as temperate) were identified with PhageLeads and BACPHLIP (≥ 95% probability only)^48,49^ . If a temperate vOTU was identified, all vOTUs within its vConTACT2 VC were also classified as temperate. Hosts were predicted for the vOTUs using iPHoP v0.9beta; a pipeline that combines RaFAH^50^, WIsH^51^, oligonucleotide frequencies^52^, PHP^53^, and BLAST ^54^. Additional host assignment was performed by comparing CRISPR matches to a compendium of rumen MAGs using PILER-CR and SpacePHARER 5-c2e680a^55,56^.

### Micro- and Macro-Diversity Statistics

Each read set was randomly down-sampled to the size of the smallest sample in which ≥1 could be detected by read mapping (Calf 3: 481,471 x 2 paired end reads) using seqtk with -s 100 (https://github.com/lh3/seqtk). Rarefied reads were mapped to the vOTUs using Bowtie 2 v2.3.4.3^57^ with --non-deterministic --maxins 2000, as described in the MetaPop paper. MetaPop was performed with --genome_detection_cutoff 75 --no_viz^58^.

### Mapping to Reference Databases

Reads were mapped separately to the INPHARED database (March 2024)^39^, a set of vOTUs produced from a virome analysis of a dairy cattle slurry tank on the same farm ^8^, NCBI viral RefSeq (March 2024)^59^, high-quality representative genomes from the Unified Human Gut Virome Catalog (https://github.com/snayfach/UHGV), the Rumen Virome Database (RVD)^17^, and the Unified Rumen Phage Catalogue (URPC)^25^ using bbmap.sh v38.84 with minid=0.90 ambiguous=best ^27^.

### Bacterial Metagenomics

Raw data was parsed for quality using fastp with its default settings^60^. The quality checked data was used as the input for bwa-mem v0.7.17^61^ against the *Bos taurus* reference genome (ARS-UCD1.3) for host sequence removal. Kraken2 v2.1.2, with minimum hit groups set to 3, was used to determine putative taxonomic assignments with the standard database^62^. The output of Kraken2 v2.1.2 was parsed through Bracken v2.8 twice, once for genus and once for species level assignments, with a read threshold of 10 to estimate abundance^63^.

### Statistics

Due to the relatively small sample sizes per group (n=3-5) and the potential non-normality of data distribution, non-parametric statistical tests were employed throughout the analysis. Kruskal-Wallis tests were used for comparisons across all four life stages, followed by Dunn’s post-hoc tests with Bonferroni correction for multiple testing when significant differences were detected. For direct comparisons between two groups (e.g., dry vs. lactating adults), Wilcoxon rank-sum tests (Mann-Whitney U) were applied. Correlations between viral and bacterial metrics were assessed using Spearman’s rank correlation coefficient. Beta-diversity analyses were performed using PERMANOVA with 999 permutations, and homogeneity of multivariate dispersions was confirmed using permutation tests (permutest). All statistical analyses were conducted in R (v4.2.1), with significance defined at p < 0.05, though marginally significant results (0.05 < p < 0.10) are noted where biologically relevant.

## Data Availability

Raw reads are available within the European Nucleotide Archive under project PRJEB52149. Virome assembly and complete viral genomes are available on FigShare at https://figshare.com/s/

## Acknowledgments

R.C was supported by a scholarship from the Medical Research Foundation National PhD Training Programme in Antimicrobial Resistance Research (MRF-145-0004-TPG-AVISO). J.H, S.H, A.M, D.S, C.D, M.J were funded by NERC AMR-EVALFARMS (NE/N019881/1). Bioinformatics analysis was carried out on infrastructure provided by MRC-CLIMB (MR/L015080/1) and CLIMB-BIG-DATA (MR/T030062/1). C.M was supported by a scholarship from the Biotechnology and Biological Sciences Research Council BBDTP (BB/M008770/12115419). R.C. and E.M.A. are funded through the Biotechnology and Biological Sciences Research Council (BBSRC) grant Bacteriophages in Gut Health BB/W015706/1. E.M.A. gratefully acknowledges the support of the BBSRC; this research was funded by the BBSRC Institute Strategic Programme Food Microbiome and Health BB/X011054/1 and its constituent projects BBS/E/F/000PR13631 and BBS/E/F/000PR13633; and by the BBSRC Institute Strategic Programme Microbes and Food Safety BB/X011011/1 and its constituent projects BBS/E/F/000PR13634, BBS/E/F/000PR13635 and BBS/E/F/000PR13636.

## Contributions

RC, AM and MJ conceived the idea. RC, CM, JR and CH collected samples and/or extracted DNA. RC, CM, AM, AMB, AP, JR and EMA analysed the data. DS, JH, MJ and AM supervised the work. RC and AM drafted the manuscript. All authors read and approved the final manuscript.

## Ethics Declarations

The authors have no competing interests to declare. Sample collection for this project was reviewed and approved by the SVMS ethics committee on the 14th of November 2017 with approval number 2132 171010.

